# Reprogramming breast cancer cells to breast cancer stem cell-*like* by the POU1F1 transcription factor

**DOI:** 10.1101/2025.04.30.651395

**Authors:** Leandro Ávila, Samuel Seoane, Sandra Rodriguez-Gonzalez, Magda Gois, Mª Efigenia Arias, David Martinez-Delgado, Noemi Lado, Tomas Garcia- Caballero, Pablo Aguiar, Roman Perez-Fernandez

## Abstract

Breast cancer stem cells (BCSCs) have been proposed as the cause of resistance to conventional treatments and as the origin of breast cancer recurrence and metastasis. However, BCSC origin is not fully understood. Using breast cancer cell lines with POU1F1 overexpression and knockdown as well as immunodeficient mice models, this study provides compelling evidence of the role of POU1F1 in reprogramming breast cancer cells into BCSC-*like* by deregulating CD24, CD44, CD133, and ALDH markers. In addition, a subpopulation from POU1F1 overexpressing MCF-7 cells with high levels of ALDH present functional changes, e.g., higher invasion, glycolytic metabolism, increased clonogenicity and mammosphere formation, high tumor-initiating capacity, and resistance to treatment. Mechanistically, these features seem to be mediated by POU1F1 activation of the IL-6/JAK2/STAT3 pathway. Our data suggests that hormone therapy-resistant Luminal A breast tumors with high levels of POU1F1 could be targeted by blocking the IL-6/JAK2/STAT3 pathway.

## INTRODUCTION

During cancer progression, cancer cells can acquire molecular and phenotypic changes, an ability known as cellular plasticity (1). Cell plasticity, also known as lineage plasticity, could be defined as the ability of a cell to reprogram and change its phenotype identity (1–3). The acquisition of epithelial-mesenchymal transition (EMT) (4–6) and cancer stem cell-*like* (CSC-*like*) states are two of the best-known cases of tumor cell plasticity, and both states have been linked (7,8). CSCs, in particular breast cancer stem cells (BCSCs), have been proposed as the cause of resistance to conventional treatments and as the origin of breast cancer recurrence and metastasis (9). Thus, metastasis could well be initiated by a subpopulation of cancer cells (also called “tumor-initiating cells”) that possess stem cell-*like* properties, being more favorable to reinitiate tumor growth at distant sites and to acquire resistance to therapy, resulting in increased aggressiveness and relapse (10–13). The most common markers for identifying BCSCs are CD24^−/low^, CD44^+/high^, CD133^+^, and ALDH^+^, which are also found in several other types of tumors (14). In addition, BCSCs have common functional characteristics, such as higher clonogenicity, formation of three-dimensional (3D) structures in cultures (called spheroids or mammospheres), and tumor-initiating capacity *in vivo* (15–16).

The POU class 1 homeobox 1 (POU1F1 or Pit1) transcription factor belongs to the **P**it1-**O**ct3/4-**U**nc86 (POU) family of transcription factors that play key roles in cell proliferation, cell lineages, and regulation of terminal differentiation (17). POU1F1 was first discovered in the pituitary gland, being responsible for cell differentiation and as a transcriptional activator during its organogenesis (18–20). However, POU1F1 is also expressed in other tissues, such as the mammary gland. In breast, POU1F1 expression in tumors is higher than in normal cells and induces cell proliferation (21,22). In addition, POU1F1 upregulates Snai1 and induces EMT, promoting tumor growth and metastasis (23). POU1F1 also influences the tumor microenvironment (TME), modifying the phenotype of fibroblasts and recruiting and differentiating normal macrophages into tumor-associated macrophages (TAMs) to promote cancer progression (24,25).

Given that POU1F1 is related to EMT-cell plasticity and metabolism modification, we intend to analyze the relationship between POU1F1 and CSC in breast cancer. Using breast cancer cell lines and immunodeficient mouse models after overexpression and knockdown of POU1F1, we evaluated changes in BCSC markers, colony growth, mammosphere formation, tumor initiation capacity, and resistance to hormone and radiotherapy. In addition, we evaluated the mechanisms by which these changes may occur.

Our results show that POU1F1 reprograms breast cancer cells to BCSC- *like* and suggests a therapeutic strategy for treating POU1F1-overexpressing breast tumors.

## RESULTS

### POU1F1 reprograms the MCF-7 cells in BCSC-*like*

To investigate the role of POU1F1 in cellular reprogramming, the luminal A subtype of human breast cancer MCF-7 cells was stably transfected with the POU1F1 overexpression vector (Supplementary Fig. 1A-B), and a gene set enrichment analysis (GSEA) was performed. We compared the transcriptomes of MCF-7 (which exhibit low endogenous POU1F1 expression) with those of MCF-7 cells overexpressing POU1F1 (hereinafter referred to as POU1F1 cells) (GSE287812). This analysis revealed significant enrichment of gene signatures associated with cellular plasticity and stem cell activation in the POU1F1 cells, suggesting that POU1F1 promotes transcriptional programs linked to cellular reprogramming (Fig. 1A–B). The volcano plot analysis of RNA-seq data in POU1F1 cells (GSE287812) reveals deregulation of gene expression related to cancer stem cell differentiation, e.g. CD44, ITGA6 (CD49f), EGFR, NT5E, CD24, EpCAM, FOXA1, and NOTCH3 (Fig. 1C). These data were further compared with data obtained from an RNA-seq in human mammary epithelial cells (HUMEC) with and without POU1F1 overexpression (GSE287732). Similar changes were not observed in the aforementioned genes related to cancer stem cell differentiation, suggesting that the effect of POU1F1 is exclusive to tumor cells (Supplementary Fig. 2).

**Fig. 1.**
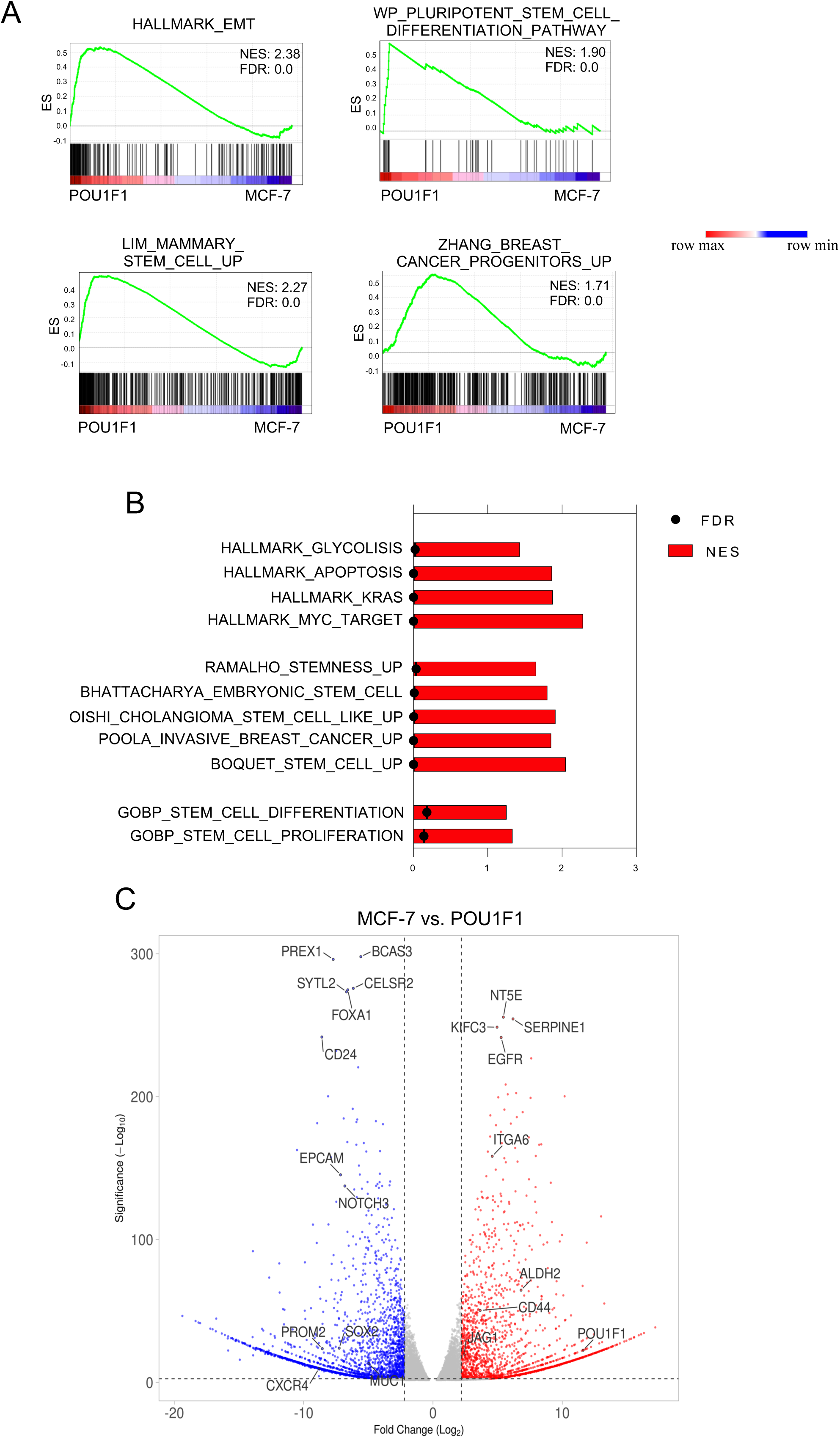
RNA-seq data of MCF-7 cells without/with POU1F1 overexpression. **A-B**. GSEA plot of enrichment in EMT, pluripotent stem cell differentiation pathway, mammary stem cell, and breast cancer progenitors geneset and dataset enrichment graph of signature associated with stem cell data by RNA-seq from POU1F1-overexpressing MCF7 cells (POU1F1) vs MCF7 cells (GSE287812). **C.** Volcano plot shows significant downregulated (blue dots) and upregulated (red dots) CSC markers in MCF-7 versus POU1F1 cells.

Subsequently, we selected well-known BCSC markers, such as CD24, CD44, CD133, and ALDH, and carried out real-time PCR in MCF-7 control cells as well as in POU1F1 cells. We found a significant increase in CD44 (P<0.001), CD133 (P<0.0001), and ALDH1A1 (P<0.01) mRNA and a decrease (P<0.0001) in the CD24 marker in POU1F1 cells (Fig. 2A). A flow cytometry was also carried out in these cells to evaluate protein expression. Double-labeling indicates a strong shift from CD24^+^/CD44^−^ markers in MCF-7 control cells to CD24^−^/CD44^+^ in POU1F1 cells. In addition, there was also an increased protein expression in CD133 and ALDH activity in POU1F1 cells (Fig. 2B). The epigenetic status of CD24, CD44, and ALDH1A1 gene promoters in POU1F1 cells show different grades of methylation, an epigenetic mark associated with activation (CD44, ALDH1A1) or suppression of transcription (CD24) (Supplementary Fig. 3). Conversely, the MDA-MB-231 cell line (triple-negative subtype, with higher endogenous levels of POU1F1 than MCF-7 cells) after POU1F1 knockdown (Supplementary Fig. 1C-E) presents a significant change in the BCSC-marker profile, i.e. higher CD44, CD133, and ALDH1A1 mRNA expression in control MDA-MB-231 cells (MDA-shC), as compared to POU1F1-knocked-down MDA-MB-231 cells (shPOU1F1) (Fig. 2C). Protein levels of CD44 and CD133 also decreased in shPOU1F1 cells, with a striking decrease in ALDH activity from 95% in control cells to 0.33% (Fig. 2D). Next, we correlated mRNA expression of POU1F1 and BCSC markers in the GSE65194 database of human breast tumor samples. Tumors were separated according to high (above the mean) or low (below the mean) CD24, CD44, CD133, and ALDH1A1 mRNA expression (n=130 tumors, Fig. 2E, upper panel). As shown in Figure 2E, lower panel, we found a significant correlation between POU1F1 and CD44 (R=0.34, P=0.002), CD133 (R=0.21, P=0.04), and ALDH1A1 (R=0.23, P=0.03) mRNA expression.

**Figure 2.**
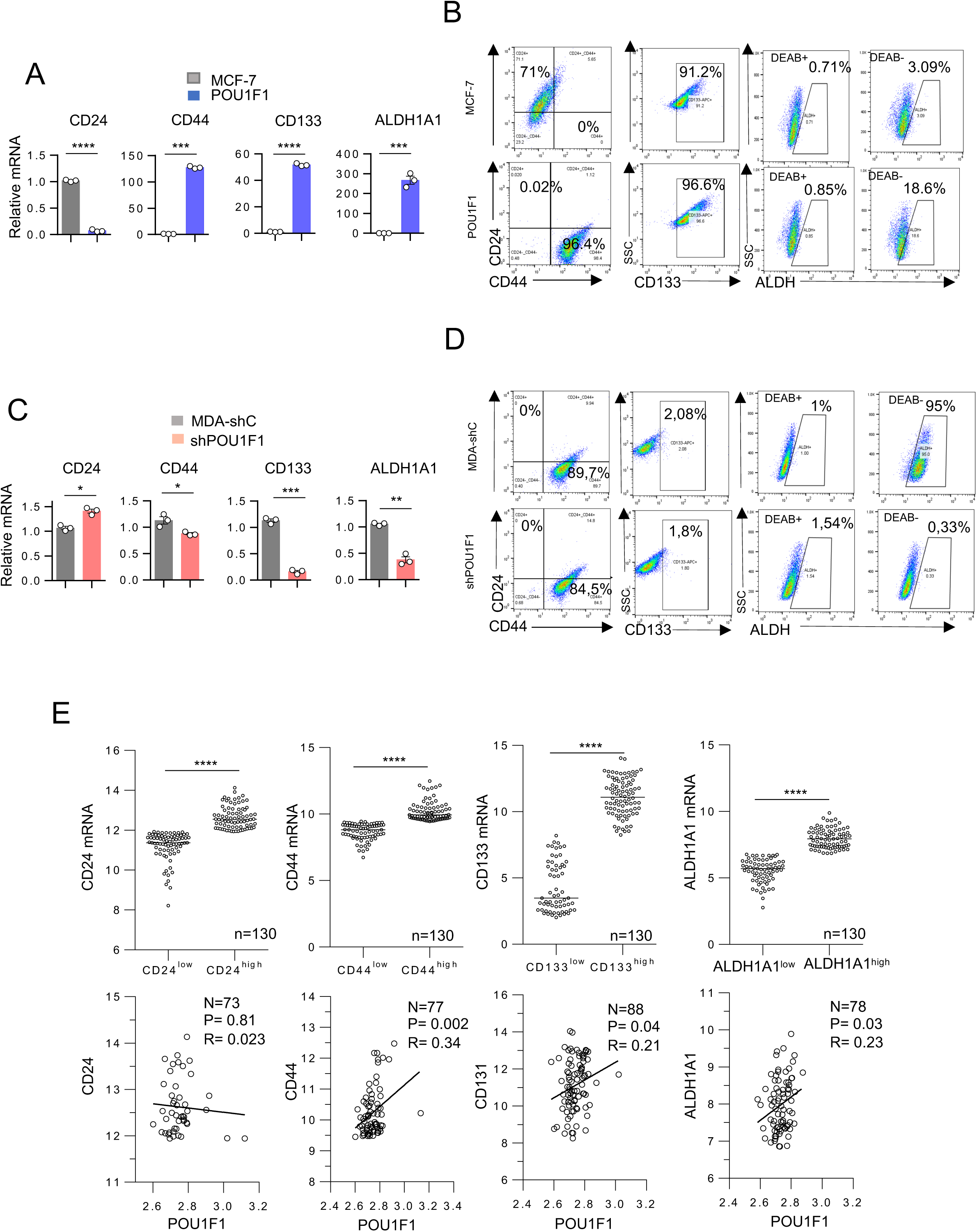
POU1F1 induces a BCSC-*like* phenotype from breast cancer cells. **A.** qPCR of CD24, CD44, CD133, and ALDH1A1 mRNA in MCF-7 and MCF-7-POU1F1 (POU1F1) cells. **B**. Flow cytometry analysis of CD24/CD44, CD133, and ALDH in MCF-7 and POU1F1 cells. Cells were double-stained with anti-CD44-PE and anti-CD24-FITC, CD133-APC, and ALDH-496 nm (DEAB was used to establish baseline fluorescence to define the ALDEFLUOR positive area). **C.** qPCR of CD24, CD44, CD133, and ALDH1A1 mRNA in control MDA-MB-231 cells (MDA-shC) and MDA-MB-231 cells with POU1F1 knock-down (shPOU1F1). **D.** Flow cytometry analysis of CD24/CD44, CD133, and ALDH in control (MDA-shC) and MDA-MB-231 cells with POU1F1 knock-down (shPOU1F1), as described in B. **E**. Dispersion plot of CD24, CD44, CD133, and ALDH1A1 mRNA levels in human breast tumors (n = 130) (GSE65194). CD24, CD44, CD133, and ALDH1A1 were classified according to their mRNA expression levels: high (above the mean) or low (below the mean). Spearman correlation analysis of POU1F1 and CD24 (n=73), CD44 (n=77), CD133 (n=88), and ALDH (n=78) mRNA expression. Results are represented as mean ± SEM. ∗p < 0.05, ∗∗p < 0.01, ∗∗∗p < 0.001, ∗∗∗∗p < 0.0001.

### A subpopulation of POU1F1 cells with high ALDH expression (POU1F1-ALDH^high^) has strong BCSC-*like* characteristics

To delve into how POU1F1-overexpressing MCF-7 cells modulate their phenotype, flow cytometry was performed to separate two populations of cells according to ALDH1A1 expression: a) POU1F1-ALDH^low^, and b) POU1F1-ALDH^high^ (Fig. 3A). As shown in Figure 3B-C, cells enriched with ALDH1A1 mRNA and protein (POU1F1-ALDH^high^) were obtained. Wound healing and invasion assays were performed to analyze the functional properties of these cells, particularly their migratory and invasive capacity. A significant increase in both migration (P<0.0001) and invasion (P<0.05) was observed in POU1F1-ALDH^high^ cells as compared to POU1F1 cells (Fig. 3D-E). POU1F1-ALDH^high^ cells metastasize significantly more in lung, brain, and liver tissues than POU1F1 cells (Fig. 3F). As previously shown (25), POU1F1 cells have a glycolytic profile because POU1F1 transcriptionally up-regulates the key lactate dehydrogenase A (LDHA) enzyme, which leads MCF-7 cell metabolism from pyruvate to lactate production. Next, the glycolytic activity of the POU1F1-ALDH^high^ subpopulation was assayed by determining extracellular acidification rate (ECAR), basal glycolysis, and compensatory glycolysis. Figure 3G-H shows that POU1F1-ALDH^high^ cells significantly increase both basal glycolysis (P<0.01) and compensatory glycolysis (P<0.05) as compared to POU1F1 cells.

**Figure 3.**
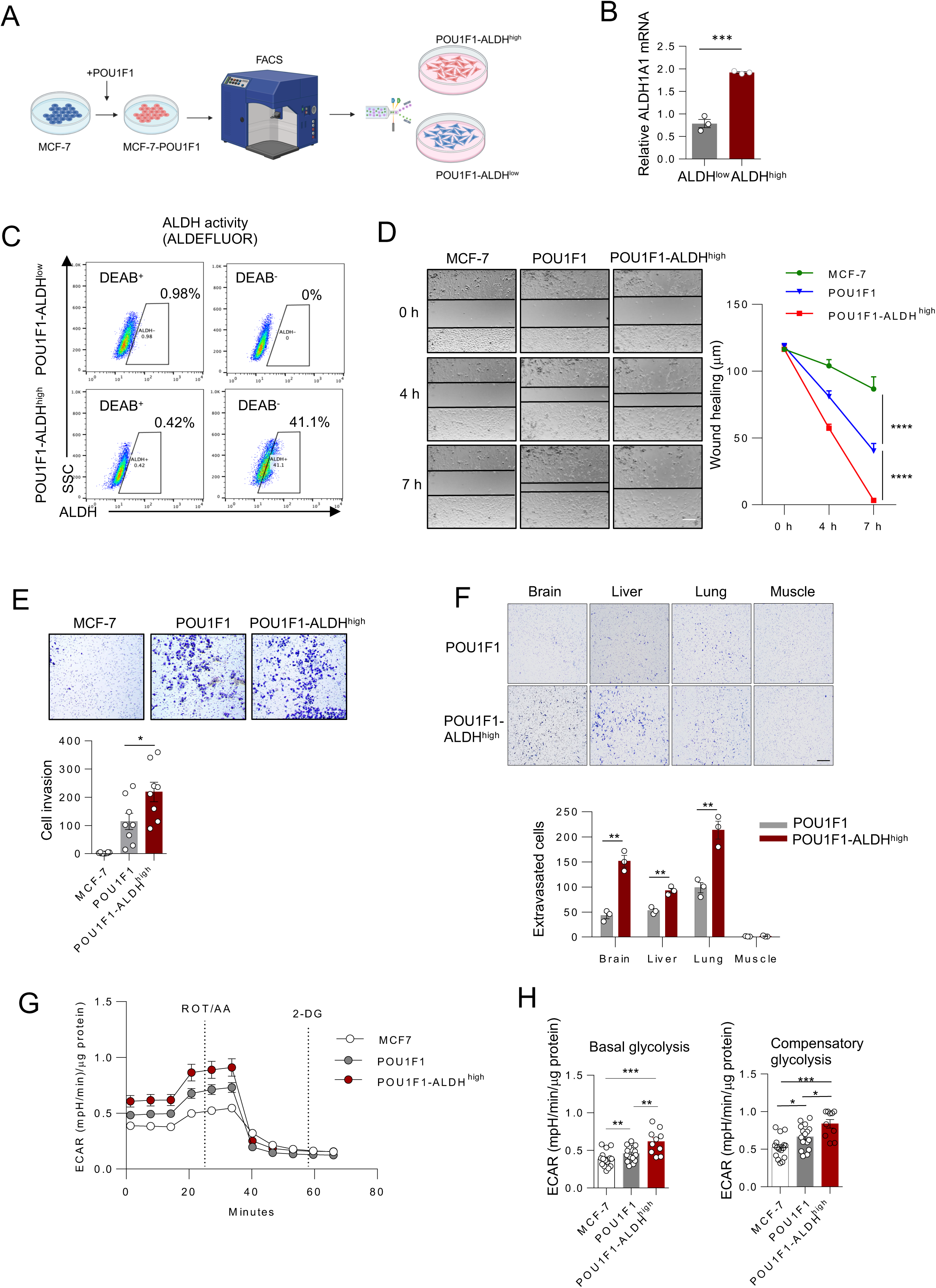
The POU1F1-ALDH^high^ subpopulation has strong BCSC-*like* characteristics. **A**. Schematic representation of isolation of the POU1F1-ALDH^high^ subpopulation. **B**. qPCR of ALDH1A1 mRNA in ALDH^low^ and ALDH^high^ cells after sorting MCF-7-POU1F1 cells. **C.** Flow cytometry analysis of ALDH activity (ALDEFLUOR assay) in POU1F1-ALDH^low^ and POU1F1-ALDH^high^ cells. **D**. Wound-healing assay of MCF-7, POU1F1, and POU1F1-ALDH^high^ cells at 0, 4, and 7 h (scale bar: 100 μm). **E**. Trans-well invasion assay and quantitative analysis in MCF-7, POU1F1, and POU1F1-ALDH^high^ cells after 24 h (scale bar: 100 μm). **F**. Trans-well invasion assay and quantitative analysis in MCF-7, POU1F1 and POU1F1-ALDH^high^ cells. Cells are plated into the upper chamber of the transwell and migrate across the matrigel and pore membrane system to the lower chamber filled with mouse tissue extract (brain, liver, lung, and muscle used as negative control). **G.** Representative ECAR Glycolytic Rate Assay profile in MCF7, POU1F1, and POU1F1-ALDH^high^ cells. **H** Quantification of basal glycolysis, and compensatory glycolysis in MCF7, POU1F1, and POU1F1-ALDH^high^ cells. Results are represented as mean ± SEM. ∗p < 0.05, ∗∗p < 0.01, ∗∗∗p < 0.001.

### Colony formation and mammosphere density are increased in POU1F1-ALDH^high^ cells

BCSCs are characterized by an increased size and number of forming colonies and the ability to form mammospheres. Thus, we studied these parameters in MCF-7, POU1F1, and POU1F1-ALDH^high^ cells. As shown in Figure 4A, high levels of ALDH significantly increase colony number (P<0.001), even though mammosphere volume is larger in POU1F1 cells than in POU1F1-ALDH^high^ cells (P<0.0001) (Fig. 4B). However, confocal microscopy indicated that the mammospheres from POU1F1-ALDH^high^ cells have a significantly (P<0.001) higher density compared to POU1F1 cells, as measured by the number of live cells (Fig. 4C). The CD49 and EpCAM markers, characteristic of progenitor luminal cells (CD49^+^) and differentiated luminal cells (EpCAM^+^), were evaluated by qPCR in these mammospheres, showing that POU1F1-ALDH^high^ cells present significantly low EpCAM (P<0.001) and high CD49 (P<0.01) mRNA expression as compared to POU1F1 cells (Fig. 4D). Conversely, POU1F1 knockdown in MDA-MB-231 (shPOU1F1) cells indicated that cells with low levels of POU1F1 have a significantly lower capacity to form colonies (P<0.0001, Fig. 4E). Moreover, mammospheres presented lower volume (P<0.0001, Fig. 4F), and lower density (P<0.01, Fig. 4G) and presented significantly high EpCAM (P<0.01) and low CD49 (P<0.001) mRNA expression as compared to MDA-shC cells (Fig. 4H).

**Figure 4.**
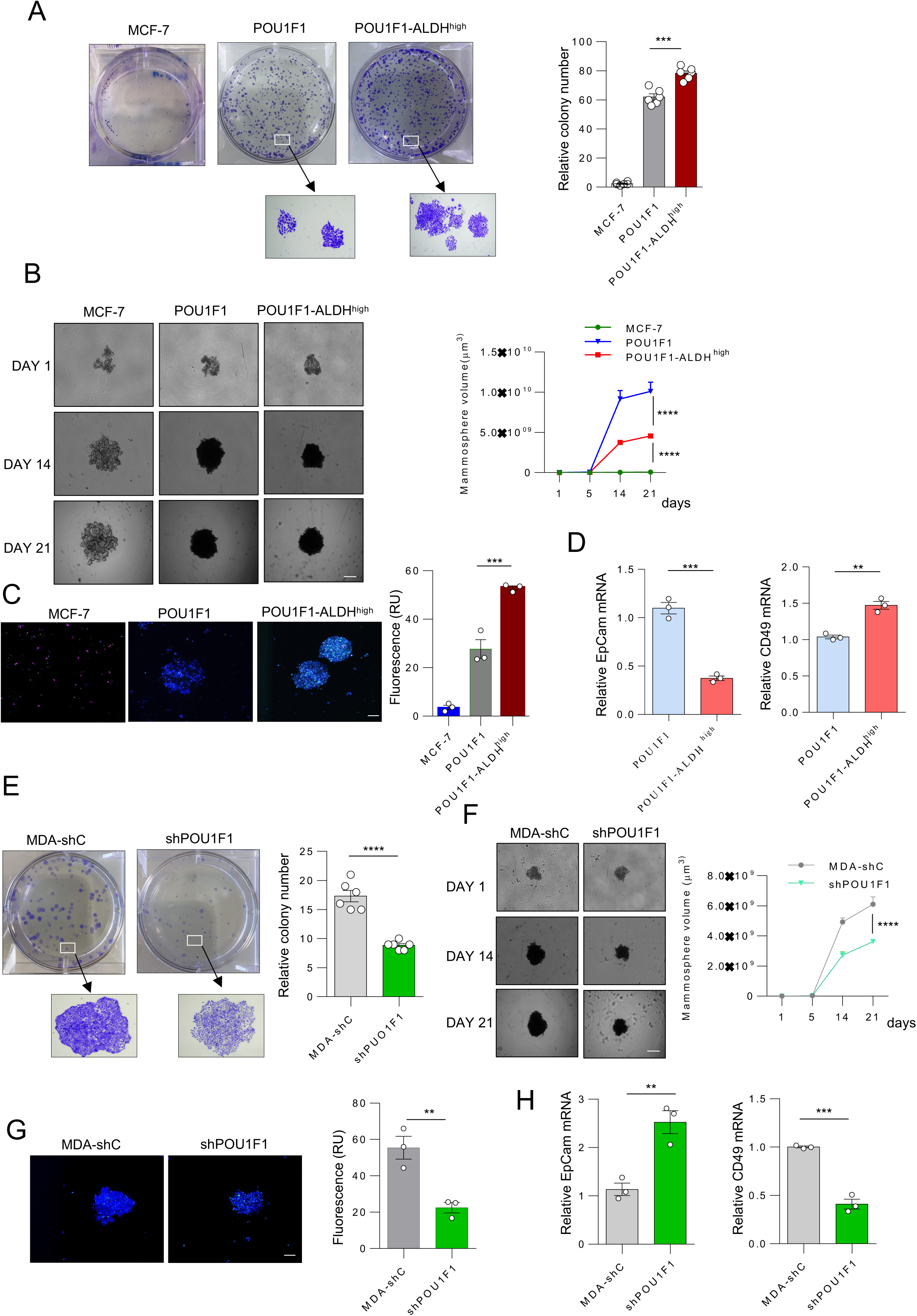
POU1F1-ALDH^high^ cells displays BCSC-*like* characteristics. **A**. Colony formation and quantitative analysis of MCF-7, POU1F1, and POU1F1-ALDH^high^ cells. **B**. Mammosphere formation and volume in MCF-7, POU1F1, and POU1F1-ALDH^high^ cells at 1, 14, and 21 days (scale bar: 200 μm). **C**. Confocal microscopy and quantification of fluorescence (live cells) at the midsection of mammospheres from MCF-7, POU1F1, and POU1F1-ALDH^high^ cells (scale bar: 200 μm). **D**. qPCR of EpCAM and CD49 mRNA in POU1F1 and POU1F1-ALDH^high^ cells. **E**. Colony formation and quantitative analysis in MDA-MB-231 control cells (MDA-shC) and MDA-MB-231 cells with knockdown of POU1F1 (shPOU1F1). **F**. Mammosphere formation and volume in MDA-MB-231 control cells (MDA-shC) and MDA-MB-231 cells with knockdown of POU1F1 (shPOU1F1) at 1, 14, and 21 days. **G**. Confocal microscopy at the midsection of mammospheres in control (MDA-shC) and POU1F1 knock-down MDA-MB-231 (shPOU1F1) cells. **H**. qPCR of EpCAM and CD49 mRNA in control (MDA-shC) and POU1F1 knock-down MDA-MB-231 (shPOU1F1) cells. Results are shown as mean ± SEM. ∗∗p < 0.01, ∗∗∗p < 0.001.

### POU1F1-ALDH^high^ cells have a greater capacity to initiate tumors and greater resistance to treatment

We next evaluated the effects of POU1F1-ALDH^high^ cells on tumor developing capacity. Immunodeficient BALB/c-nu female mice were injected in the mammary fat pad with either 500 or 5,000 cells of MCF-7, POU1F1, and POU1F1-ALDH^high^, and a tumor-initiating capacity (TIC) assay was performed. Of the mice injected with POU1F1-ALDH^high^ cells, 3 out of the 8 mice injected with 500 cells and 5 out of the 7 mice injected with 5,000 cells developed tumors after 6 weeks; compared with 2 out of 8 mice and 0 out of 7 mice in the POU1F1-injected group. TIC frequency was significantly higher (2,769) in POU1F1-ALDH^high^-injected mice compared to POU1F1 (19,249) (P=0.006) and MCF-7 (36,443) (P=0.001) injected mice (Fig. 5A). Figures 5B-C show tumor luminescence and examples of HE staining with mitosis indicated by white arrows. As shown in Supplementary Fig. 4, tumors from MCF-7 injected cells are smaller than those injected with POU1F1 cells.

**Figure 5.**
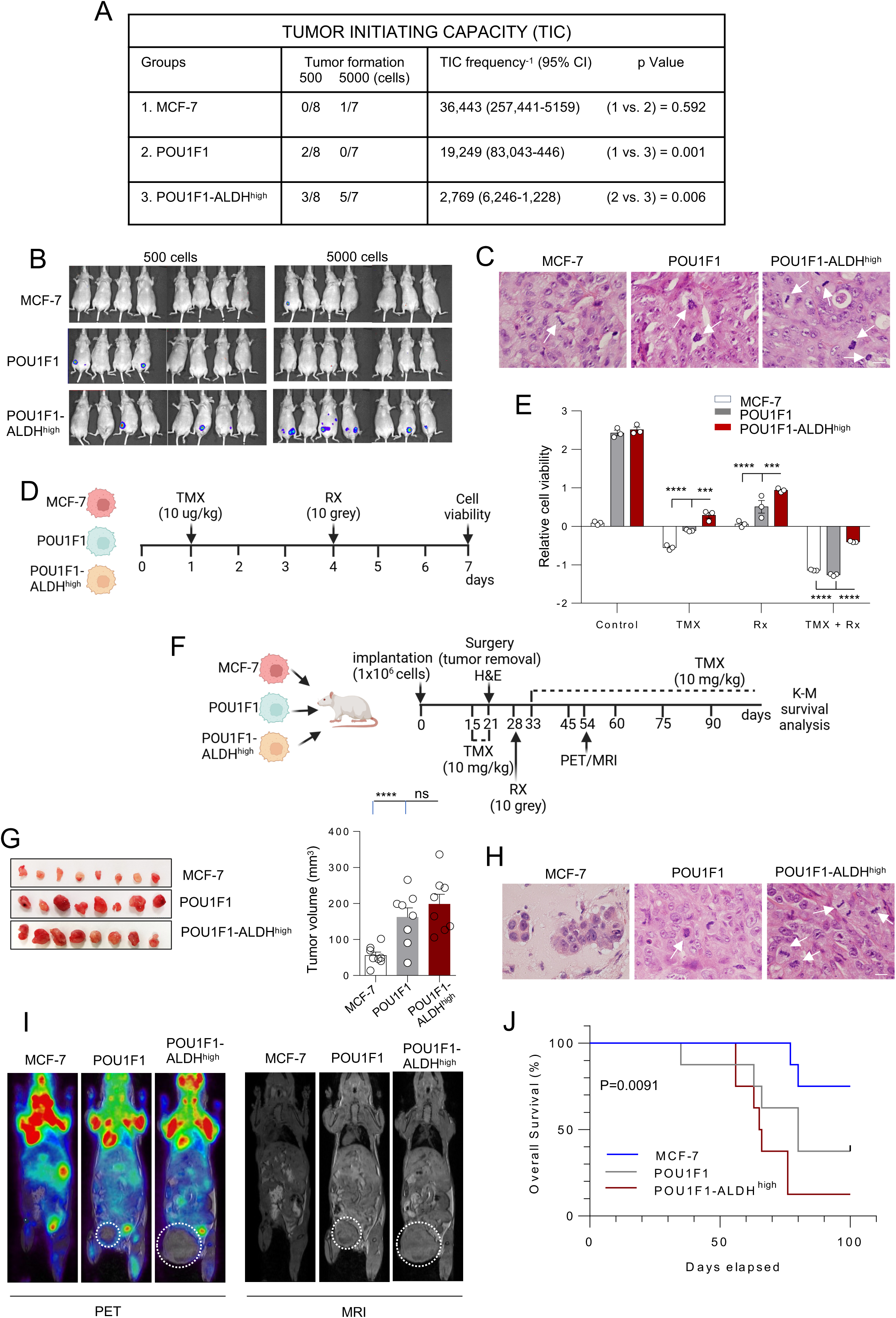
Evaluation of TIC and resistance to treatment. **A**. Tumor-initiating capacity (TIC) of MCF-7, POU1F1, and POU1F1-ALDH^high^ cells implanted into the fourth mammary fat pad of mice (5×10^2^ cells, n=8, and 5×10^3^ cells, n=7). Tumors were monitored for 6 weeks to assess tumorigenicity. **B**. Luminescence of tumors (IVIS images) was recorded at 6 weeks. **C.** Example of HE staining in tumors. White arrows indicate cell mitosis (low in MCF7 cells, intermediate in POU1F1, and high in POU1F1-ALDH^high^ cells). **D**. Experimental design to evaluate hormone- and radiotherapy resistance in MCF-7, POU1F1, and POU1F1-ALDH^high^ cells (TMX: tamoxifen; RX: radiotherapy). **E**. Relative cell viability in MCF-7, POU1F1, and POU1F1-ALDH^high^ cells after hormone therapy, radiotherapy, and both. **F**. Experimental design to evaluate hormone- and radiotherapy resistance *in vivo* after xenografting of MCF-7, POU1F1, and POU1F1-ALDH^high^ cells in mice (n=8 per group). **G**. Tumors were excised 21 days post-implantation of cells into the mammary fat pad, and tumor volume was calculated. **H**. Examples of H&E staining in tumors. White arrows indicate mitosis. **I**. [^18^F] FDG PET/MRI assessed glucose uptake and tumor growth on day 54 in two mice per group. The dotted circle highlights the tumor in PET images. **J**. Kaplan-Meier overall survival curves. Results are presented as mean ± SEM. ∗∗∗p < 0.001, ∗∗∗∗p < 0.0001, ns: not significant.

We developed an *in vitro* study using hormone therapy and radiotherapy to evaluate tumor cell resistance to treatment. Tamoxifen (TMX), a selective estrogen receptor modulator, clinically used for the treatment of hormone receptor-positive Luminal A subtype of breast tumors (such as the MCF-7 cell line), was administered to MCF-7, POU1F1, and POU1F1-ALDH^high^ cells, and 72 h later, a radiation dose was given. Cell viability was then analyzed (Fig. 5D). Higher cell viability, i.e., higher resistance to treatment (TMX, radiotherapy, and TMX + radiotherapy) was observed in POU1F1-ALDH^high^ cells compared to the other groups (Fig. 5E). We also tested the resistance to treatment *in vivo* (Fig. 5F). After treatment with TMX on day 15 for 6 days, tumor was surgically excised under anesthesia. Figure 5G shows no significant difference in tumor volume between mice injected with POU1F1-ALDH^high^ cells and those injected with POU1F1 cells. Tumor H&E staining suggests a higher number of cell mitoses in the POU1F1-ALDH^high^ group, although these data were not statistically calculated (Fig. 5H). After a single dose of radiotherapy on the excised tumor area on day 28, mice were treated again with TMX from day 33 until death, and overall survival was assessed by Kaplan-Meier analysis. In addition, on day 54, glucose uptake was analyzed in two mice per group using [^18^F] FDG PET/MRI (Fig. 5I). Control MCF-7 mice showed no evidence of tumors in either the MRI or PET images. POU1F1 mice showed an active tumor in one mouse, which was small in the morphological images (indicated by a dotted circle) but with significant metabolic activity at the tumor edges. POU1F1-ALDH^high^ mice showed large, spherical tumors in the MRI images, with metabolic activity primarily located at the tumor edges, while the central region of the tumor appears inactive, likely indicating necrotic tissue (Fig.5I and Supplementary Fig. 5). Finally, the Kaplan-Meier survival analysis shows a clear difference in overall survival among the three groups over time (P=0.009) (Fig. 5J). MCF-7 had the highest survival rate, with no significant decline until later time points. POU1F1-ALDH^high^ demonstrated the poorest survival, with a rapid and steep decline in survival rates observed early in the study period compared with POU1F1 cells.

### The POU1F1 activates the IL-6/JAK2/STAT3 pathway

To elucidate the possible mechanisms involved in the reprogramming of breast cancer cells to BCSC-*like* by POU1F1, bioinformatic analyses for inflammatory and cytokine processes involved in CSC activity were carried out in an RNA-seq of MCF-7 and POU1F1 cells (GSE287812). POU1F1 overexpression induced a clear enrichment of hallmarks of inflammatory response and Interleukin-6/Janus kinase 2/signal transducer and activator of transcription 3 (IL-6/JAK2/STAT3) signaling pathway signatures (NES=2.18, FDR=0.00, and NES=1.88, FDR=0.004, respectively) (Fig. 6A). The IL-6/JAK2/STAT3 pathway has been involved in CSC marker regulation in several tumor types (26–28). In addition to IL-6, other members of the IL-6 cytokine family include IL-11, oncostatin M (OSM), leukemia inhibitory factor (LIF), cardiotrophin 1 (CT-1), ciliary neurotrophic factor (CNTF), cardiotrophin-like cytokine factor 1 (CLCF1), IL-27, IL-35, and IL-39 (29). A heatmap from our RNA-seq data indicates a significant up-regulation in CNTF, IL-6, IL-11, CLCF1, LIF, and CT-1 following POU1F1 overexpression (Fig. 6B). Indeed, Volcano plot analysis revealed that multiple cytokines were upregulated after POU1F1 overexpression, including IL-6 and IL-11 with a fold change > 3.5 (Supplementary Fig. 6A). Both qPCR and ELISA analyses confirm that POU1F1 significant (P<0.0001) increases IL-6 mRNA expression, and IL-6 protein in culture medium (Fig. 6C). Also, IL-11, CLCF1, LIF, and CT-1 mRNA significantly increased after POU1F1 although not CTNF mRNA (Fig. 6D). Indeed, POU1F1 induces phosphorylation of proteins involved in the JAK2/STAT3 signaling pathway, such as STAT3, ERK1/2, and AKT (Fig. 6E). To study whether POU1F1-reprogramming of cancer cells could be mediated by ALDH through the IL-6/JAK2/STAT3 pathway, POU1F1 cells were treated with two Janus Kinase inhibitors (JKI), Ruxolitinib and AG490 (Supplementary Fig. 6B). A significant decrease in ALDH1A1 mRNA and protein activity was found (Fig. 6F-G). In addition, MCF-7 cells treated either with conditioned medium (CM) from POU1F1 cells or the IL-6 ligand show a significant (P<0.001 and P<0.01, respectively) ALDH1A1 mRNA increase (Fig. 6H-I). On the contrary, POU1F1 cells treated with the IL-6 ligand and an anti-IL-6 receptor antibody (Tocilizumab-TCZ, used clinically to block the IL-6R pathway) indicated a significant (P<0.01) decrease in ALDH1A1 mRNA (Fig. 6I). Furthermore, TCZ administration to POU1F1 cells significantly reduces colony and mammosphere formation (P<0.01) as compared to control cells (Fig. 6J-K).

**Figure 6.**
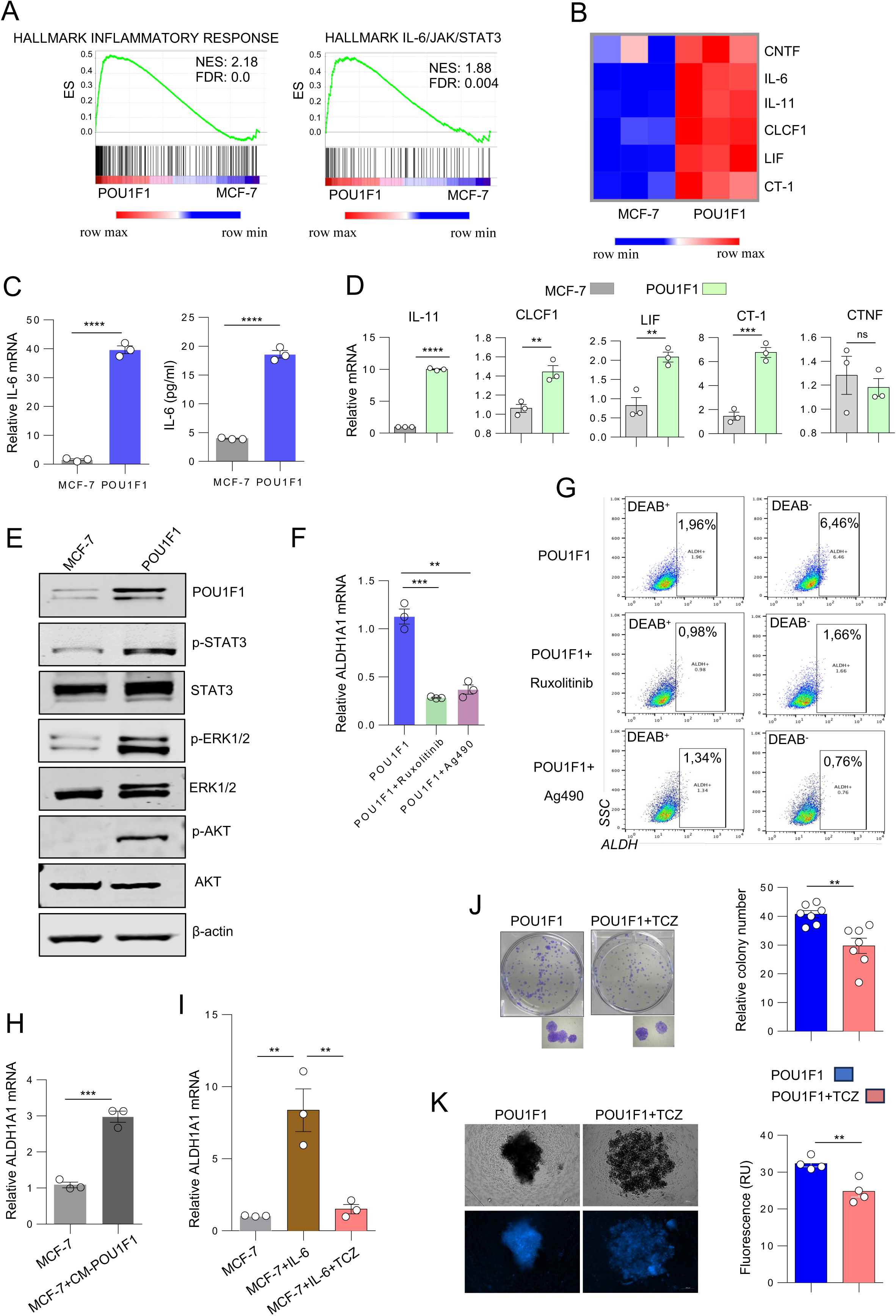
POU1F1 regulates the IL-6/JAK2/STAT3 pathway. **A**. GSEA plot of enrichment in inflammatory response and IL-6/JAK2/STAT3 pathway of the RNA-seq data from POU1F1-overexpressing MCF-7 (POU1F1) cells vs. MCF-7 cells. **B.** Heat map of some IL-6 family of interleukins with and without POU1F1 overexpression in MCF-7 cells. **C**. qPCR of IL-6 mRNA and ELISA of IL-6 protein in culture medium after 72 h. **D**. qPCR of IL-11, CLCF1, LIF, CT-1, and CNTF mRNA expression in MCF-7 and POU1F1 cells. **E.** Immunoblots of POU1F1, phosphorylated (p) STAT3 (p-STAT3), STAT3, p-ERK1/2, ERK1/2, p-AKT, AKT, β-actin in control MCF-7 and POU1F1 cells. **F**. qPCR of ALDH1A1 mRNA in POU1F1 cells before and after treatment with Ruxolitinib and Ag490. **G**. Flow cytometry analysis of ALDH activity (ALDEFLUOR assay) in POU1F1 cells before and after treatment with R uxolitinib and Ag490. **H**. qPCR of ALDH1A1 mRNA expression in MCF-7 untreated cells before and after treatment with conditioned medium (CM) from a 72-h culture of POU1F1-overexpressing MCF-7 cells. **I**. qPCR of ALDH1A1 mRNA in MCF-7 control cells before and after treatment with IL-6, and with IL-6+Tocilizumab (TCZ, anti-IL-6 receptor antibody). **J**. Colony formation in POU1F1 cells before and after treatment with Tocilizumab (TCZ). **K.** Mammosphere formation in POU1F1 cells before and after treatment with Tocilizumab (TCZ). Live cells were detected by fluorescence at the midsection of mammospheres (scale bar: 200 μm) Results are presented as mean ± SEM. ∗∗p < 0.01, ∗∗∗p < 0.001, ∗∗∗∗p < 0.0001.

## DISCUSSION

POU1F1 is a well-known transcription factor involved in embryonic and adult pituitary development that has also been analyzed for its role in breast cancer (23). This study provides compelling evidence of the role of POU1F1 in reprogramming breast cancer cells into BCSC-*like*. Our data demonstrate that POU1F1 induces a BCSC-*like* phenotype in breast tumor cells by deregulating markers such as CD24, CD44, CD133, and ALDH. These phenotypic modifications correlate with functional changes, i.e., higher clonogenicity, increased mammosphere formation, and increased glycolytic metabolism. In addition, using *in vivo* immunodeficient mice models, we found that a subpopulation of ALDH^high^ cells obtained from the POU1F1 overexpressing MCF-7 cells possess high tumor-initiating capacity, high resistance to hormone and radiotherapy, and reduced overall survival. Mechanistically, these actions seem to be mediated by POU1F1 through activating the IL-6/JAK2/STAT3 pathway.

The overexpression of POU1F1 in the luminal A subtype of the breast cancer MCF-7 cell line (with low POU1F1 endogenous levels) induces a phenotypic shift from CD24^⁺^/CD44^⁻^ and ALDH^low^ into CD24^⁻^/CD44^⁺^ and ALDH^high^, characteristics of BCSCs, as first described by Al-Haji et al. (15). Conversely, in the triple-negative MDA-MB-231 cell line (with higher endogenous POU1F1 levels), blockade of POU1F1 results in decreased expression of CD44, CD133 and a striking drop in ALDH activity. This shift in BCSC markers correlated with POU1F1 mRNA expression levels in human breast tumor databases, independently of the tumor subtype. However, POU1F1 overexpression does not induce phenotypic changes in normal human mammary epithelial cells, suggesting that the effects of POU1F1 are restricted to the specific tumor context. The subpopulation isolated from POU1F1 cells and identified as POU1F1-ALDH^high^ cells exhibits the highest functional capacities, including enhanced migration and invasion compared to other subpopulations. These findings highlight its distinctive potential and are consistent with previously described characteristics of cells with similar phenotypes (30,31). Increased basal glycolysis may correspond to the high energy demands of BCSCs during the growth and proliferation process (32). Furthermore, these cells show a marked increase in compensatory glycolysis, indicating metabolic flexibility and the ability to adapt to metabolic stress, such as hypoxia or oxidative stress induced by free radicals.

A higher cell density was observed in mammospheres from POU1F1-ALDH^high^ cells, and the CD49f^+^/EpCAM^−^ phenotype acquired by these cells could be associated with the ability to generate different structures within the mammary gland including basal, luminal progenitor, and mature cell types (33), suggesting a potential role in differentiation and organization of the tumor microenvironment (TME). Moreover, this phenotype has been identified as a marker associated with a high metastatic potential in tumor cells and an increased risk of disease recurrence (34).

The BCSC-*like*, specifically the POU1F1-ALDH^high^ subpopulation, also shows higher tumor-initiating capacity (TIC). It has been described that TIC creates a microenvironment that spatially favors tumor progression and drug resistance (35). Specifically in breast cancer, the ALDH1A1 isoform in TICs was described as promoting tumor progression through immune system modulation (36). Also, the results revealed that POU1F1 significantly enhances resistance to conventional therapies, a process amplified by elevated ALDH levels. High expression of ALDH, driven by POU1F1, appears to impart tumor cells with critical reprogramming capabilities, allowing them to adopt a phenotype that promotes therapeutic evasion. This reprogramming leads to increased tumor aggressiveness, higher relapse rates, and reduced overall survival, underscoring the pivotal role of POU1F1 in cancer progression and treatment resistance. ALDH is defined as a molecular target to study drug resistance in cancer stem cells, and the downregulation of ALDH1A1 increases the sensitivity of tumor cells (37). CSCs overexpressing ALDH showed lower ROS levels than differentiated cancer cells, probably due to increased antioxidant enzymes (38). In breast cancer, ALDH has been associated with resistance to other treatments, such as PARP inhibitors (39).

Mechanistically, we described a connection between POU1F1 and the IL-6/JAK2/STAT3 pathway. This pathway has been associated with pluripotency, therapy resistance, and metastasis (40). After POU1F1 overexpression, there was a significant increase in the RNA-seq concerning inflammatory response and the IL-6/JAK2/STAT3 pathway signature. RNA-seq data were confirmed by qPCR, showing increased IL-6 mRNA and protein levels in the culture medium and other members of the IL-6 ligand family. The increased activity of this POU1F1-mediated pathway may at least partially explain the role of ALDH in BCSC reprogramming. The specific activation of STAT3 has been linked to the promotion of c-MYC, KLF4, and SOX9 expression, which support the self-renewal of BCSC-ALDH^high^, as well as mechanisms of chemoresistance (41,42). The abnormal activation of JAK2/STAT3 signaling driven by the IL-6 ligands family has also been positively correlated with EMT and metastasis in lung tumors (43). In addition, recent findings highlight the differential regulation of the EMT process by STAT3 phosphorylation at specific residues, depending on the molecular subtype of CSCs (44). Furthermore, STAT3 activation triggers downstream proteins such as AKT and ERK1/2, which are well-characterized for their roles in cellular metabolism, proliferation, metastasis, and CSC maintenance mechanisms (45). Tyrosine kinase inhibitors (TKI) and different drugs against the ligands of the IL-6 family have been used to prevent the activation of the signaling pathway at various checkpoints, thus decreasing expression of stemness markers, i.e., CD44, CD133, ALDH, and EpCAM, and mammosphere formation in TNBC tumors (40). Our data agree with these data and demonstrate the functional effects of TKI administration and IL-6R blockade on ALDH activity and colony and mammosphere formation, suggesting these drugs as a possible treatment in tumors with POU1F1-overexpression.

In summary, our study reveals a novel role for POU1F1 in reprogramming breast cancer cells into BCSC-*like* and the role of ALDH^high^ subpopulations in aggressive traits. We showed that POU1F1-overexpressing cells mediated ALDH regulation by the IL-6/JAK2/STAT3 signaling pathway. Our results highlight a possible use of tyrosine kinase inhibitors (TKIs) or anti-IL-6R antibodies targeted against BCSCs in breast tumors with POU1F1 overexpression by disrupting the POU1F1/IL-6/JAK2/STAT3/ALDH1A1 axis and thus preventing tumor recurrence.

## MATERIALS AND METHODS

### Cell culture and drugs

The human breast adenocarcinoma MCF-7 and MDA-MB-231 cells were obtained from ATCC-LGC (Barcelona, Spain). Cell lines were grown in DMEM (Sigma, St Louis, USA) supplemented with 10% FBS, 100 U/ml penicillin-streptomycin (Thermo Fisher Scientific, Waltham, USA), at 37 °C in 5% CO_2_. The human mammary epithelial cell (HUMEC-TERT) was obtained from BIOCAT (Heidelberg, Germany) and grow in DMEM F12 supplemented with 5% horse serum, 0.5% penicillin/streptomycin, 0.1mM glutamax, and 12.5 ng/ml ascorbic acid (Thermo Fisher Scientific), 0.5µg/ml hydrocortisone and 1% fetal bovine pituitary extract (STEMCELL Technologies, Vancouver, Canada), 10 µg/ml insulin, 20 ng/ml hEGF, and 100 ng/ml cholera toxin (Thermo Fisher Scientific) at 37 °C in a 5% CO_2_. Cell lines were tested and authenticated according to microscopic morphology, growth curve analysis, and mycoplasma detection according to the ECACC cell line verification test recommendations. Tamoxifen was administered to cells at 10 μg/ml every other day and 10 mg/kg for 28 days *in vivo*. This study used two Janus kinase inhibitors, Ruxolotinib (a dose of 498 nM) and Ag490 (a dose of 15 nM) (STEMCELL Technologies), and an anti-IL-6 receptor (IL-6R) antibody, Tocilizumab (a dose of 30 μg/ml, Thermo Fisher Scientific). IL-6 (Merck, Darmstad, Germany) was used a dose of 100 ng/ml.

### Plasmid and transfections

Stable overexpression of POU1F1 in the MCF-7 cells (POU1F1 cells) was achieved through lentiviral infection of a POU1F1 overexpression plasmid (pLV-puro-EF1A>hPOU1F1/FLAG) obtained from Vector Builder (Chicago, USA), which contains an ORF clone of the human POU1F1 gene. The empty vector (pLV-puro-EF1A) was used as a control. POU1F1 blockade was performed using a pool of 3 target-specific lentiviral vector plasmids, each encoding 19-25 nt (plus hairpin) shRNAs to knockdown the POU1F1 gene expression (Santacruz Biotechnology, Heidelberg, Germany). A pool of 3 scrambled shRNA sequences was used as a control.

### Cell viability, migration, and invasion assays

Cell viability/proliferation was measured using an MTT assay (Thermo Fisher Scientific). Cells were seeded in 6-well plates at a density of 5×10⁴ cells per well. After 72 h, absorbance was measured at 590 nm using an LB 940 Mithras multi-plate reader (Berthold Technologies, Shangai, China).

Wound-healing assays were performed using an insert system in 35-well mm plates (IBIDI, Martinsried, Germany). Cells (3×10⁴) were seeded in each insert in DMEM. After 24 h, the inserts were removed, and the plates were placed under the Widefield Leica DMI 6000B microscope (Leica) to conduct a 12 h time-lapse analysis. Invasion assay was performed in 24-well plates containing inserts with 8 μm pores (Corning, New York, USA). At the top of the inserts, a three-dimensional structure resembling the extracellular matrix was created using Geltrex (Thermo Fisher Scientific), and 2.5×10⁴ cells were seeded in serum-free DMEM. Culture medium supplemented with 20% FBS or protein extract (200 mg/ml) from mouse-specific tissues (brain, liver, lung, and muscle) was added to the bottom of the inserts. After 24 h of incubation, cells that migrated through the pores onto the bottom of the insert were fixed in 90% cold methanol and stained with crystal violet (Sigma). The total number of invading cells was determined by counting the cells on the lower surface of the insert using an inverted microscope (Olympus IX73, Olympus, Allentown, USA).

### Colony and spheroid/mammosphere formation assays

Colony formation assays were performed in 6-well plates with 100 cells/well. Colonies grew for 7 days, fixed with 90% methanol for 30 min at RT, and stained with crystal violet solution (Sigma) for 30 min. Colonies were counted directly on the plate. For three-dimensional (3D) mammosphere formation, 500 cells/well were seeded on low-attachment Nunclon Sphera plates (Thermo Fisher Scientific). Images were captured on days 1, 14, and 21. Mammosphere volume was calculated using the formula: π/6 × length × width^2^. In addition, mammospheres were stained for 1 h with a triple staining protocol using Hoechst, caspase 3/7, and propidium iodide to identify live, apoptotic, and dead cells, respectively. Mammosphere density was analyzed by laser scanning confocal microscopy (Leica Stellaris) and Cell Discoverer 7 (Zeiss, Oberkochen, Germany) measuring the fluorescence of live cells.

### RNA isolation and qPCR

Total RNA for massive sequencing (RNA-seq) and qPCR was isolated using the commercial DNA, RNA, and protein purification kit (Macherey-Nagel, Düren, Germany). cDNA was synthesized with M-MLV-RT (Invitrogen, Thermo Fisher Scientific), and reactions of quantitative real-time PCR were done using SYBR Green PCR Master Mix (Thermo Fisher Scientific). The samples were denatured at 95°C for 10 min, annealed at 55°C for 15 sec, and extended at 72°C for 40 sec for a total of 40 cycles on Applied Biosystems™ QuantStudio™ qPCR Systems (Thermo Fisher Scientific). qPCR primers used in this study:

Human *pou1f1* forward: 5’-TCCTGACCACACCTTGAGTC-3’

Human *pou1f1* reverse: 3’-CTTTTCCGCCTGAGTTCCTG-5’

Human *18s* forward: 5’-GTAACCGCTTGAACCCCATT-3’

Human *18s* reverse: 3’-CCATCCAATCGCTAGTAGCG-5’

Human *gapdh* forward: 5’-GTAACCCGTTGAACCCCATT-3’

Human *gapdh* reverse: 3’-CCATCCAATCGCTAGTAGCG-5’

Human *cd24* forward: 5’-ACCCACGCAGATTTATTTCCA-3’

Human *cd24* reverse: 3’-ACCACGAAGAGACTGGCTGT-5’

Human *cd44* forward: 5’-AAGGTGGAGCAAACACAACC-3’

Human *cd44* reverse: 3’-AGCTTTTTCTTCTGCCCACA-5’

Human *cd133* forward: 5’-TTCTATGCTGTGTCCTGGGG-3’

Human *cd133* reverse: 3’-GGGCCCATTTTCCTTCTGTC-5’

Human *epcam* forward: 5’-CTTTAAGGCCAAGCAGTGCA-3’

Human *epcam* reverse: 3’-CCAGTAGGTTCTCACTCGCT-5’

Human cd49f forward: 5’-AGAATTGACCTCCGCCAGAA-3’

Human cd49f reverse: 3’-TTTCAGCTTCAAGTGTGCCC-5’

Human *aldh1a1* forward: 5’-AAAGAAGCTGCCCGGGAAAAG-3’

Human *aldh1a1* reverse: 3’-CCCCATGGTGTGCAAATTCA-5’

Human *il-6* forward: 5’-CTTCAGGCCAAGTTCAGGAG-3’

Human *il-6* reverse: 3’-AGTGG ATCGTGGTCGTCTTC-5’

### Western blot, ELISA, immunofluorescence (IF), and histology

Proteins were extracted using a RIPA lysis buffer (Thermo Fischer Scientific) supplemented with protease and phosphatase inhibitors. Cell extracts (30 μg) were denatured at 95°C, subjected to SDS-PAGE, and transferred to PVDF membranes (Merck Millipore, Burlington, USA). After blocking with Intercept Protein-Free Blocking buffer (LI-COR Biotechnology, Lincoln, USA), membranes were incubated overnight at 4°C with anti-POU1F1 antibody (Sigma), anti-GPDH, and anti-β-actin (Santacruz), anti-STAT3, anti-pSTAT3, anti-ERK1/2, anti-pERK1/2, anti-AKT, and anti-pAKT (Cell Signalling, Danvers, USA) antibodies for their expression and phosphorylation. A fluorescent secondary antibody was then applied, and detection was performed using the Li-Cor Odyssey fluorescence system at 700 and 800 nm wavelengths. ELISA assay to determine IL-6 protein content released to culture medium was carried out using a human IL-6 ELISA kit as the manufacturer’s instructions (Proteintech, Planegg-Martinsried, Germany).

IF assays were performed on glass coverslips pretreated with poly-L-lysine for 1 h at RT. A total of 2×10⁴ cells were seeded on the coverslips, and 24 h later, cells were fixed in cold 96% ethanol, permeabilized with PBS+0.3% Triton+0.1 M glycine for 10 min at RT and blocked with 1% BSA in PBST for 30 minutes. Primary antibodies were incubated with 0.1% BSA in PBST overnight at 4°C. After addition of the secondary antibody in 1% BSA, coverslips were mounted using DePex mounting medium Gurr (VWR, Barcelona, Spain) containing DAPI for nuclear staining. IF was analyzed using a Leica DM4B digital upright fluorescence microscope (Leica, Wetzlar, Germany). For histology, samples were fixed in 10% formaldehyde for 24 h and embedded in paraffin blocks. Sections of 4 μm were cut with a microtome and stained using a standard H&E procedure. Sections were studied in a BX51 microscope with a DP70 digital camera (Olympus).

### Flow cytometry, ALDEFLUOR assay, and cell sorting

Flow cytometry for CD24, CD44, and CD133 protein expression was carried out after labeling cells with the anti-CD24, anti-CD133, and anti-CD44 antibodies (BD) and analyzed using the Accuri Ruo Special Ryder System BD cytometer (BD) and/or the CytoFlex S cytometer (Beckman Coulter). To measure aldehyde dehydrogenase (ALDH) enzymatic activity, the ALDEFLUOR™ kit (STEMCELL Technologies) was used. Cell sorting was performed with the S3e Cell Sorter (BIO-RAD) to select POU1F1 cells with high ALDH activity (POU1F1-ALDH^high^). Data analyses were performed with FlowJo 10.0.

### Seahorse extracellular flux assays

Oxygen consumption rate (OCR) and extracellular acidification rate (ECAR) were determined using a Seahorse Bioscience XF96 Extracellular Flux Analyzer (Seahorse Bioscience, Billerica, USA). Cells (15×10^3^) were seeded into XFp microplates. After 24 h, the culture medium was changed to XF basal medium containing 10 mM glucose, 1 mM pyruvate, and 2 mM Glutamine (7.4 pH). The plate was incubated at 37°C in a non-CO_2_ incubator for 60 min before extracellular flux analysis. After four baseline measurements, the following respiration modulators were sequentially added to the wells: oligomycin (Oligo, 1.5 µM), carbonyl cyanide-4 (trifluoromethoxy) phenylhydrazone (FCCP, 1 µM), and rotenone/antimycin A (Rot/AA, 0.5 µM) for mitochondrial function assessment; or Rot/AA followed by 2-deoxyglucose (2-DG, 50 mM) for glycolytic function evaluation. Results were normalized to protein content for accurate analysis.

### Tumor-initiation analyses (limiting dilution experiments) and resistance to hormone- and radiotherapy

All animal studies were approved by the University of Santiago de Compostela Ethics Committee for Animal Experiments. For tumor-initiation capacity (TIC), eight-week-old female immunodeficient mice NRj-Foxn1^nu/nu^ (Janvier Labs, Le Genest St Isle, France) were used. Three types of cells were employed: MCF7, POU1F1 (MCF-7 cells with stable overexpression of POU1F1), and the POU1F1-ALDH^high^ cells (with high ALDH expression after sorting POU1F1 cells). Subcutaneous tumors were generated in the fourth mammary gland by inoculating two cell dilutions (500 cells, n=8 mice, and 5000 cells, n=7 mice). Cells were previously transfected with a pCDNA3.1-Hyg-CMV-fireflyLuciferase vector to monitor tumor initiation and growth. After luciferin injection (150 mg/kg), tumor growth was monitored by luminescence using the In Vivo Imaging System (IVIS, Caliper Life Sciences, Alameda, USA). An intensity map was obtained using the Living Image software (Caliper Life Sciences). The software uses a color-based scale to represent the intensity of each pixel (from blue representing low to red representing high). The TIC frequency was calculated using the Limiting Dilution Analysis software (https://bioinf.wehi.edu.au/software/elda/). Finally, tumors were removed at 8 weeks, and H&E staining was performed.

Female mice (age-matched, 8 weeks) with immunodeficiency Athymic Nude-Foxn1^nu^ (ENVIGO, Indianapolis, USA) were used for xenograft studies to evaluate the resistance to treatments. Orthotopic tumors were generated in mice by inoculation into the mammary fat pad of: a) 1×10^6^ MCF7 cells (n=8), b) 1×10^6^ POU1F1-overexpressing MCF-7 cells (POU1F1, n=8), and c) 1×10^6^ ALDH^high^ cells sorted from POU1F1-overexpressing MCF-7 cells (POU1F1-ALDH^high^, n=8) in 0.2 ml of DMEM (without FBS) and Matrigel (50:50, BD Biosciences). Tumor growth was monitored externally using a digital caliper. On day 15, mice were treated with Tamoxifen (TMX, 10 mg/kg) for 6 days - until day 21 - when tumors were surgically removed under isoflurane anesthesia (Alvira, Barcelona, Spain). Tumor volume was calculated using the formula: π/6 × length × width^2^. H&E staining of tumors was performed. Seven days later - day 28 - the mice were irradiated with a single dose of 10 gray in the tumor area, and on day 33, TMX was again administered daily (10 mg/kg) until death or sacrifice to analyze overall survival with the Kaplan-Meier method. Mice were sacrificed when primary tumors reached a volume higher than 1500 mm^3^ or when mice lost at least 20% of their total body weight.

### Positron Emission Tomography/Magnetic Resonance Imaging (PET/MRI)

PET/MRI studies using ^18^F-Fluoro-2-deoxy-2-D-glucose ([^18^F] FDG) were performed on day 54 after tumor cell injection in 2 mice per group to study tumor metabolism. All animals were fasted for 12 h before the radiotracer injection. Fused PET and MRI images were acquired using a Bruker BioSpec 3 T PET/MRI scanner (bore diameter 17 cm) equipped with actively shielded gradients (450 – 900 mT/m). After an overnight fast, 6.30 ± 0.38 MBq of ^18^F-FDG was injected into the tail vein of each animal. PET/MRI static acquisitions consisting of 10 min PET scan and 16 min MRI scan were performed. All PET images were reconstructed using the MLEM (maximum likelihood expectation maximization) 0.5 mm algorithm with 18 iterations, including scatter, randoms, and decay correction. The image pixel size was 0.5×0.5×0.5 mm3, with a FOV of 90×90×150 mm3. For MRI, a pilot scan was acquired using the Bruker Localizer protocol, with an acquisition time of 32 s. Then, an axial study was conducted using a FISP (Fast Imaging with Steady-state free Precession) sequence to cover the whole body with an Echo Time (ET) = 2.8 s, Repetition Time (RT) = 5.7 s, Averages (NA) = 5, 30.2 × 30.2 × 50 mm FOV and a matrix size of 120 x 120 × 100. Finally, animals were returned to their cages with free access to food and water. PET/MRI images were analyzed using AMIDE software. Initially, MRI images were utilized to identify the tumor region. Once the tumor regions were identified, a qualitative assessment of the [^18^F] FDG PET images was performed to evaluate tumor metabolism by visually inspecting the intensity and distribution of radiotracer uptake. To ensure a fair and consistent qualitative assessment, the color scale was uniformly adjusted across all PET images, maintaining the same scaling parameters during visualization to detect regions with increased metabolic activity.

### RNA sequencing (RNA-seq) analysis

RNA-seq was performed by Novogene (Cambridge, UK) on the Illumina NovaSeq6000 platform. High-quality reads were aligned to the human reference genome (Ensembl GRCh38, version 108) using the STAR software (v2.7.6). Once aligned, the reads were assigned to their corresponding transcripts using the htseq-count program (v0.13.5) with the “intersection-nonempty” setting to ensure that each read was mapped to a single gene. Gene expression analysis was performed using DESeq2 (v1.32.0) in R (v4.2.2), and the data matrices were enriched with additional information using BiomaRT (v2.52.0) in R. This enrichment included details such as official gene names, chromosomal location, transcription direction, and other relevant attributes. Other RNAseq data used in this study were obtained from the NCBI Gene Expression Omnibus (GEO) database (https://www.ncbi.nlm.nih.gov/geo/). Enrichment analysis of transcriptional profiles was performed with the GSEA program (Gene Set Enrichment Analysis; v4.3.2). Transcriptional profiles were compared using different gene collections and parameters: weighted, enrichment statistics, and T-Test gene ranking metric. After the analysis, enrichment plots were obtained, including the NES (Normalized Enrichment Score) and FDR (False Discovery Rate) values.

### Statistical Analysis

Values are expressed as mean ± SEM. Means were compared using Student’s t-test or analysis of variance (ANOVA) with Tukey-Kramer multiple comparison tests. The Mann-Whitney U test was used for nonparametric data with 95% confidence intervals. Correlations between mRNA expression levels were calculated using Pearson’s correlation coefficient. P-values below 0.05 were considered statistically significant. Kaplan-Meier overall survival curves were analyzed using the Log-rank (Mantel-Cox) test. Significance levels are represented as *=P<0.05, **=P<0.01, ***=P<0.001, and ****=P<0.0001 compared to the control. GraphPad Prism program was used for graph generation and statistical analyses.

## DATA AVAILABILITY

Raw and processed data for sequencing studies are available from NCBI’s Gene Expression Omnibus. They are accessible through GEO Series accession numbers GSE287812 (MCF-7 cells) and GSE287732 (human mammary epithelial cells, HUMEC) with and without POU1F1 overexpression (POU1F1) (RNA-seq). DNA methylation analyses GSE295731. All other data will be available from the authors upon request.

## AUTHOR CONTRIBUTIONS

RP-F conceptualized the study. RP-F and LAC designed the study and coordinated the experimental planning. LAC performed data and statistical analysis. LAC, MEA, and SSR were responsible for the mouse experiments. LAC, SSR, SRG, and MG were responsible for *in vitro* experiments. DMD was responsible for RNA-seq analysis. PA and NL executed PET/MRI. TG-C is responsible for H&E and tissue analysis. RPF and LAC wrote the manuscript. All authors edited and approved the manuscript.

## ETHICS APPROVAL

All animal procedures were approved by the University of Santiago de Compostela Ethics Committee (code 15012/2020/013).

## ADDITIONAL INFORMATION

### Supplementary information

Supplementary information is available on the Nature Communications website

**Supplementary Table 1.** Antibodies used.

| **Antibodies** | **Used for** | **Source** | **Dilution** |
| --- | --- | --- | --- |
| POU1F1 | WB & IF | Sigma. Ref. 089K4863 | 1:5000 |
| CD24 | FC | BD. Ref. 555428 | 1:100 |
| CD133 | FC | BD. Ref. 566596 | 5 mg/ml |
| CD44 | FC | BD. Ref. 555478 | 1:1000 |
| GPDH | WB | Sc. Ref. 32233 | 1:5000 |
| β-actin | WB | Sc. Ref. 47778 | 1:5000 |
| p-STAT3 | WB | CS. Ref. 34911 | 1:400 |
| STAT3 | WB | CS. Ref. 9132 | 1:1000 |
| p-ERK1/2 | WB | CS. Ref. 9101 | 1:1000 |
| ERK1/2 | WB | CS. Ref. 9102 | 1:1000 |
| p-AKT | WB | CS. Ref. 9271 | 1:1000 |
| AKT | WB | CS. Ref. 9272 | 1:1000 |

WB: Western blot; IF: immunofluorescence; FC: Flow cytometry; Sc: Santa Cruz; CS: Cell Signaling.

**Supplementary Figure 1.**
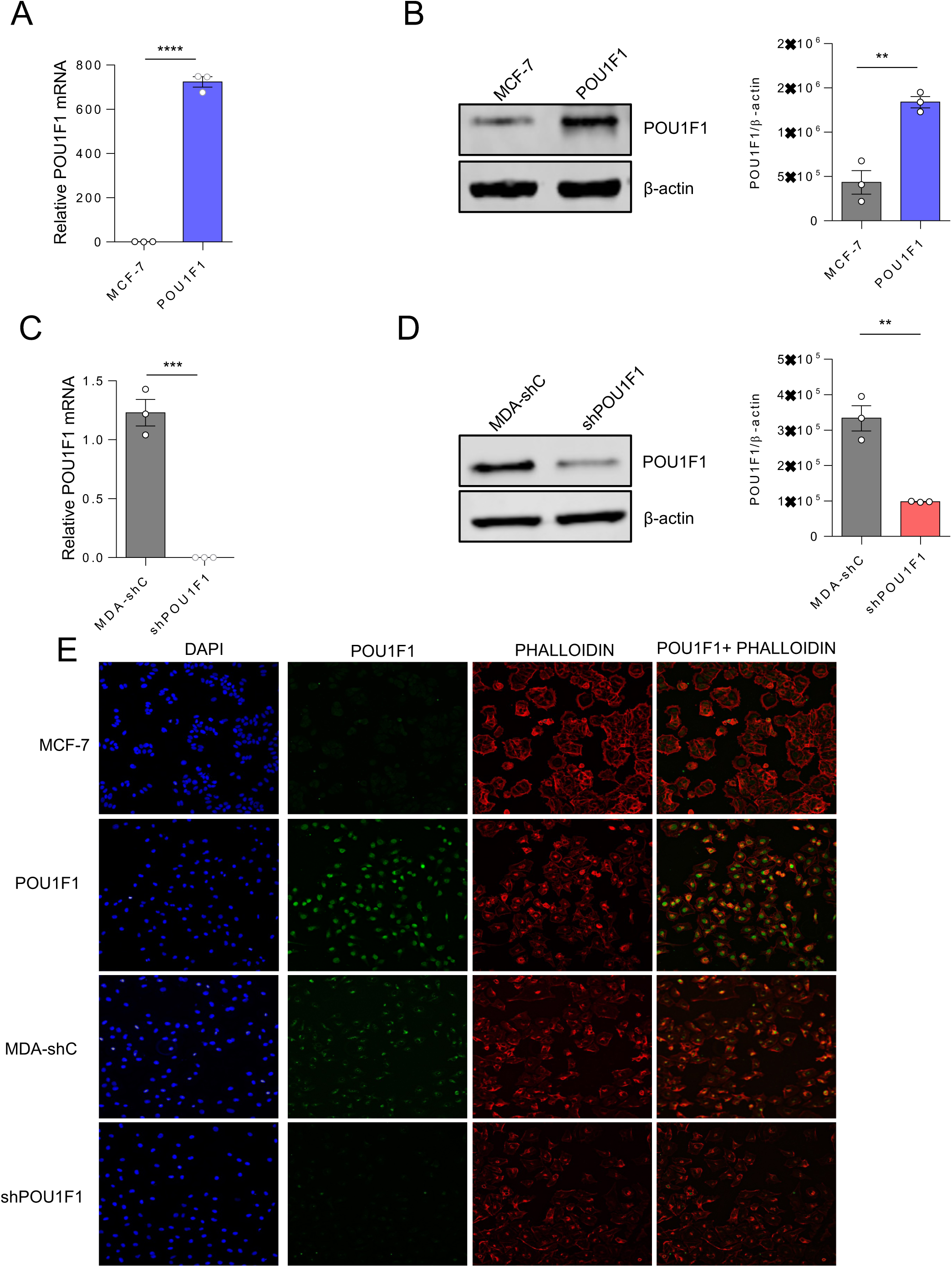
Overexpression and knockdown of POU1F1 in MCF-7 and MDA-MB-231 cells. A-B. POU1F1 mRNA and protein expression in control (MCF-7) and in POU1F1 stably transfected (POU1F1) MCF-7 cells. C-D. POU1F1 mRNA and protein expression in control MDA-MB-231 cells (MDA-shC) and after POU1F1 knockdown (shPOU1F1). E. Immunofluorescence of MCF-7, POU1F1, MDA-shC, and shPOU1F1 cells. Cells were stained with DAPI (stains the nuclei), anti-POU1F1 (stain in nuclei), and phalloidin (stains actin filaments).

**Supplementary Figure 2.**
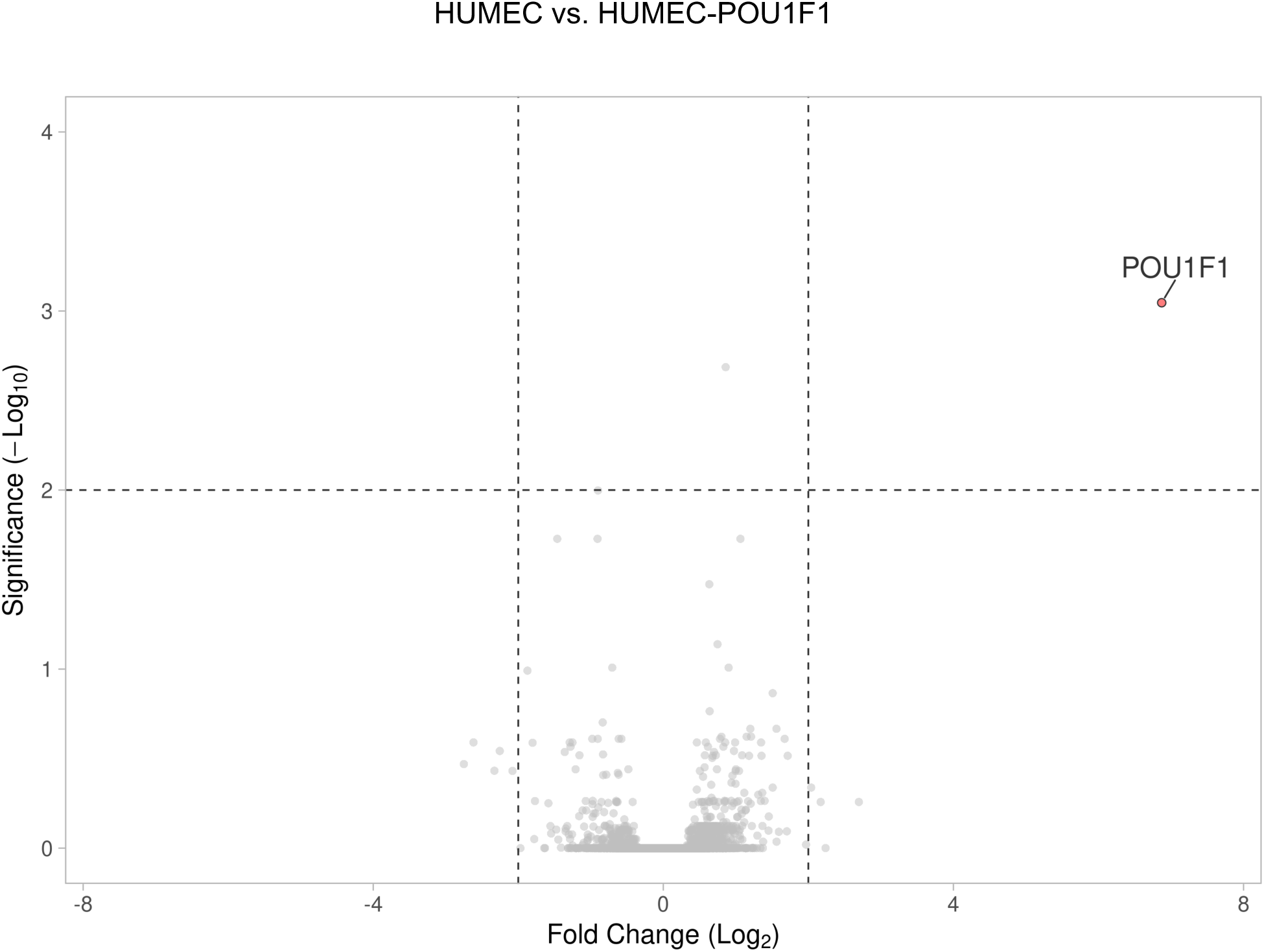
Volcano plot in HUMEC vs. HUMEC-POU1F1 cells (GSE287732).

**Supplementary Figure 3.**
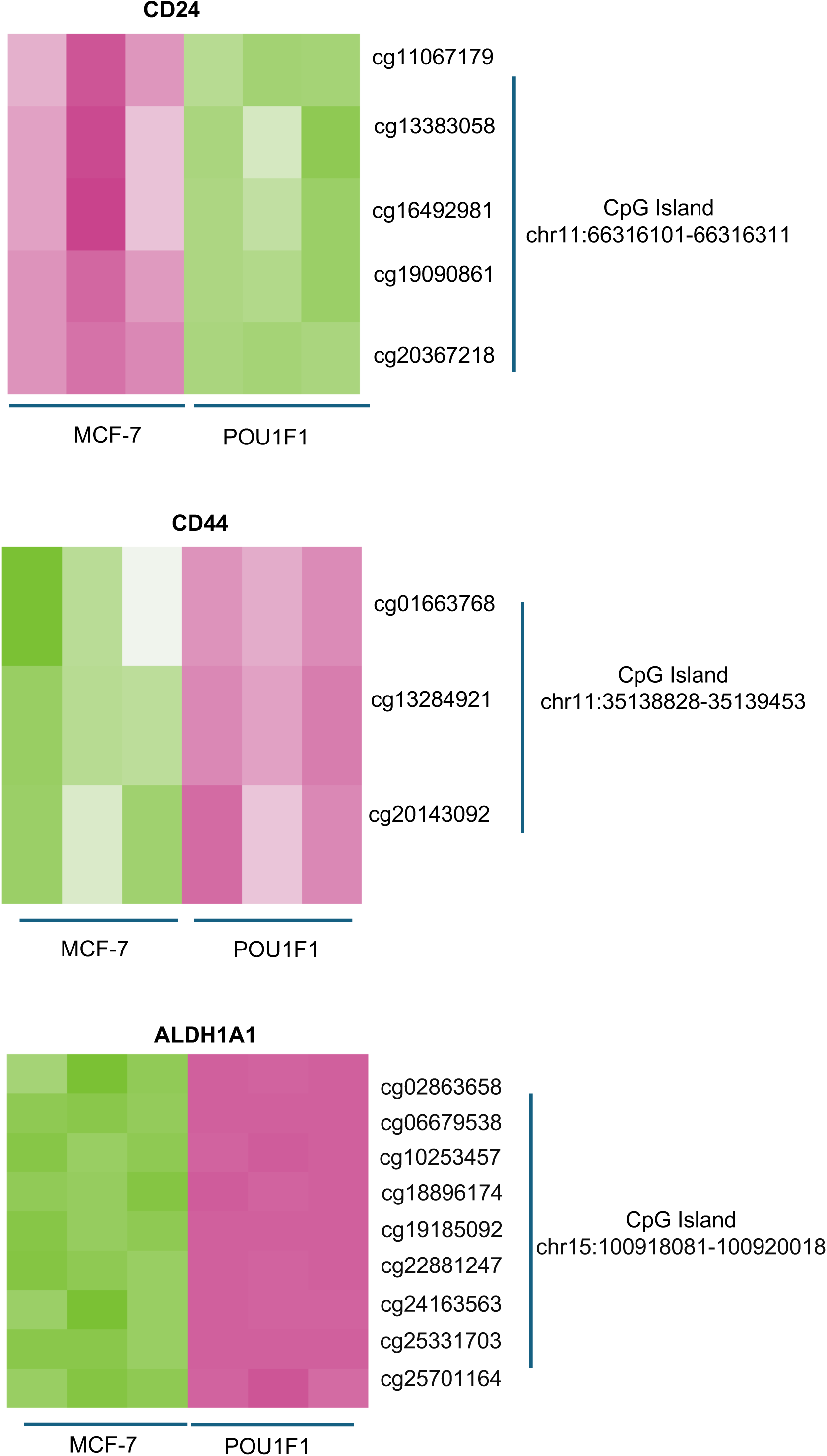
Heat-map of methylation at the CD24, CD44, and ALDH1A1 genes in MCF-7 cells before and after POU1F1 overexpression (GSE295731 ).

**Supplementary Figure 4.**
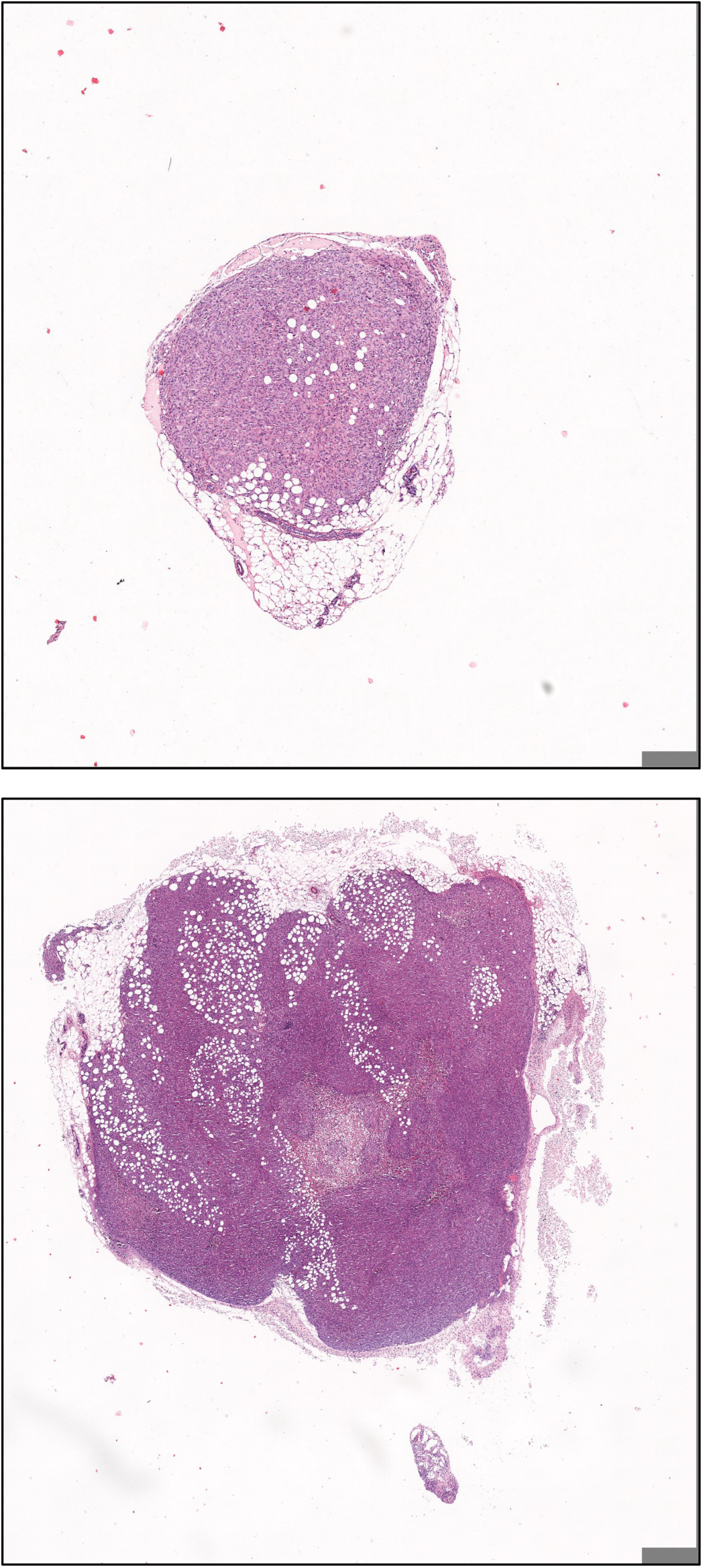
Example of H&E of tumors from mice injected with MCF-7 cells (upper) and POU1F1 cells. Note that both are at the same scale.

**Supplementary Figure 5.**
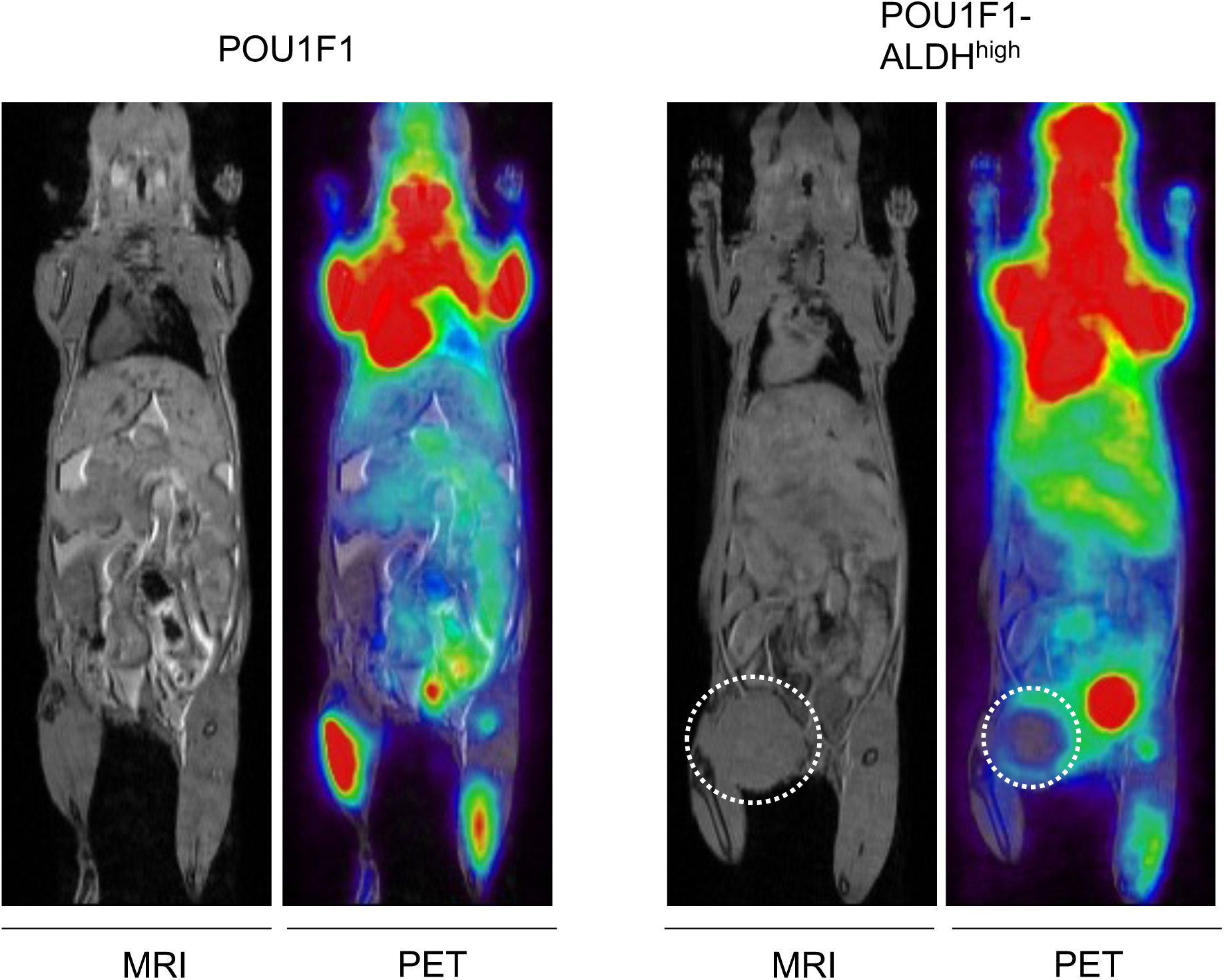
[^18^F] FDG PET/MRI assesses glucose uptake and tumor growth on day 54 in one mouse from POU1F1 and POU1F1-ALDH^high^ group. The dotted circle highlights the tumor in MRI/PET images from the POU1F1-ALDH^high^ group. In the POU1F1 group, no visible tumors are present.

**Supplementary Figure 6.**
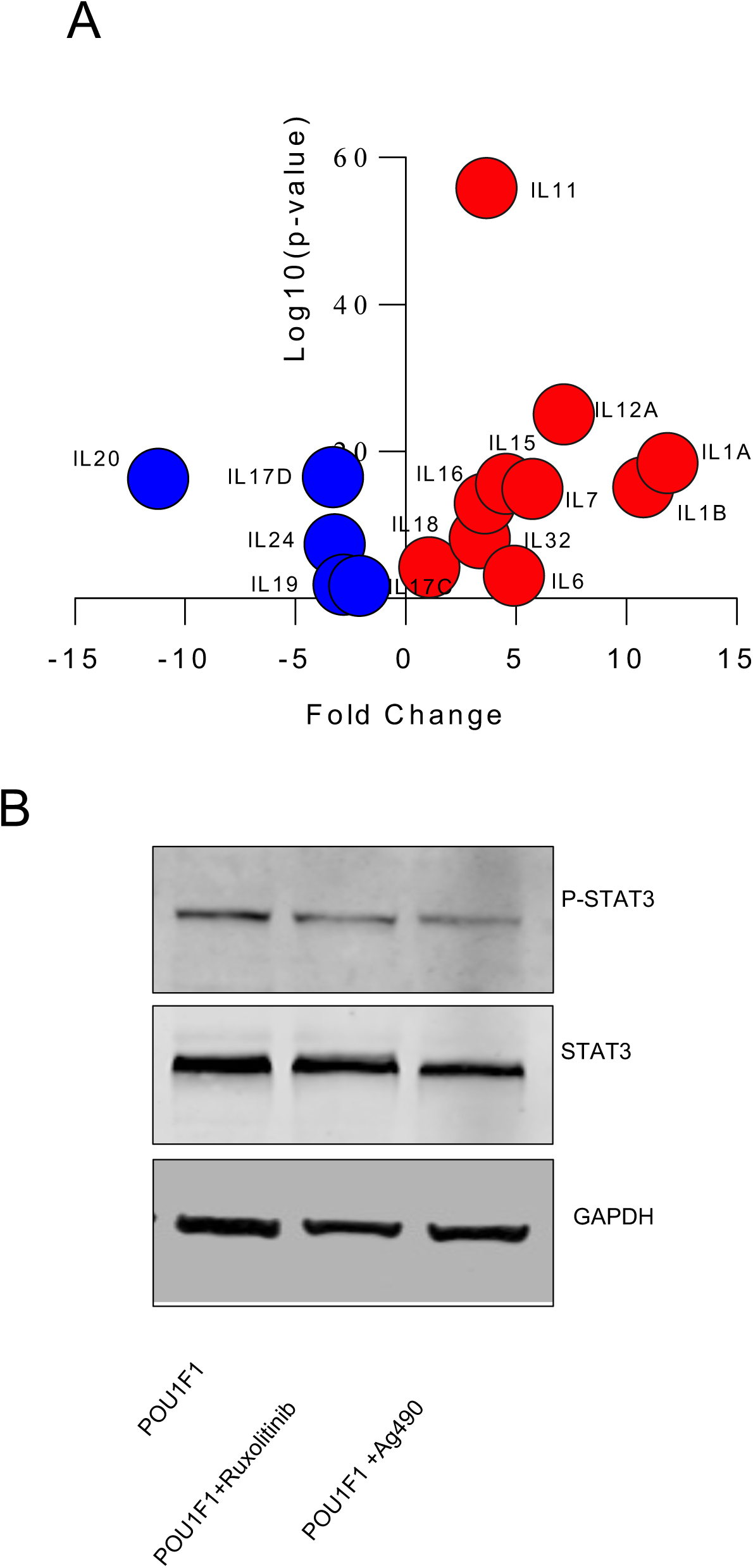
**A**. Volcano plot showing downregulated (blue dots) and upregulated (red dots) interleukins mRNA in POU1F1 vs. MCF7 cells. **B**. Immunoblots of pSTAT3 and STAT3 after treatment of POU1F1 cells with the Janus kinase inhibitors Ruxolitinib (0.428 μM) and Ag490 (12 μM)

